# Single-Cell Analysis of 5-ALA Intraoperative Labeling Specificity for Glioblastoma

**DOI:** 10.1101/2022.12.17.520870

**Authors:** Zhouzerui Liu, Angeliki Mela, Julia Furnari, Michael G. Argenziano, Corina Kotidis, Colin P. Sperring, Nelson Humala, Jeffrey N. Bruce, Peter Canoll, Peter A. Sims

## Abstract

Glioblastoma (GBM) is the most common and aggressive malignant primary brain tumor, and surgical resection is a key part of the standard-of-care. In fluorescence-guided surgery (FGS), fluorophores are used to differentiate tumor tissue from surrounding normal brain. The heme synthesis pathway converts 5-aminolevulinic acid (5-ALA), a fluorogenic substrate used for FGS, to fluorescent protoporphyrin IX (PpIX). The resulting fluorescence is thought to be specific to transformed glioma cells, but this specificity has not been examined at single-cell level. We performed paired single-cell imaging and RNA sequencing of individual cells (SCOPE-seq2) on human GBM surgical specimens with visible PpIX fluorescence from patients who received 5-ALA prior to surgery. SCOPE-seq2 allows us to simultaneously image PpIX fluorescence and unambiguously identify transformed glioma cells from single-cell RNA-seq (scRNA-seq). We observed that 5-ALA treatment results in labeling that is not specific to transformed tumor cells. In cell culture, we further demonstrated that untransformed cells can be labeled by 5-ALA directly or by PpIX secreted from surrounding transformed cells. In acute slice cultures from mouse glioma models, we showed that 5-ALA preferentially labels GBM tumor tissue over non-neoplastic brain tissue, and that this contrast is not due to blood-brain-barrier disruption. Taken together, our findings support the use of 5-ALA as an indicator of GBM tissue, but not as a specific marker of transformed glioma cells.

## Introduction

Glioblastoma (GBM) is the most common and aggressive malignant brain tumor in adults with a median survival time of <15 months (1, 2). The standard treatment for GBM is surgical resection followed by chemotherapy and radiotherapy (2). The surgical extent of resection (EOR) is associated with longer overall survival in patients with GBM (3, 4), but complete resection is impossible due to the diffusely infiltrative nature of GBM and the potential for damage to normal brain tissue. Fluorescence-guided surgery (FGS) uses fluorophores to help differentiate between normal and malignant tumor tissue in real-time during surgery to facilitate safe, maximal EOR (5, 6). 5-aminolevulinic acid (5-ALA) is the only Food and Drug Administration (FDA) approved agent for glioma FGS (7). It is a fluorogenic, heme synthesis precursor and can be metabolized to fluorescent molecule protoporphyrin IX (PpIX) by cells (8). 5-ALA has high positive predictive value (PPV) for malignant glioma tumor tissue and improves the rate of complete resection (7, 9).

The metabolic status of a cell could affect the intracellular conversion of 5-ALA to PpIX (8, 10-14). Thus, unlike simpler FGS agents such as fluorescein, 5-ALA does not rely exclusively on structural features of tumors, like blood-brain-barrier (BBB) disruption. Rather, 5-ALA is thought to directly and specifically label glioma cells. This labeling specificity could be particularly important for defining the infiltrative margins of glioma in FGS (15-17). However, although previous studies have correlated 5-ALA labeling with histologic features of glioma (18, 19), the labeling specificity of 5-ALA has not been characterized at the single-cell level. Also, false-positive 5-ALA labeling has been observed in some tissues with inflammation or reactive astrocytes (15, 20). Furthermore, due to logistical challenges with administration and intraoperative imaging, high cost, phototoxicity risk (21), and the existence of alternatives like fluorescein (22-24), more detailed characterization of 5-ALA’s advantage in labeling specificity is needed.

Flow cytometry can potentially measure PpIX fluorescence and phenotypes of individual cells (25). However, transformed glioma cells are highly heterogeneous and can resemble multiple neural lineages in the brain (26, 27). Thus, there are no specific and pervasive epitopes for unambiguously identifying transformed glioma cells across patients. SCOPE-seq is an integrated single cell imaging and RNA sequencing method (28, 29). This method enables simultaneous measurement of cell PpIX fluorescence by imaging and unambiguous identification of transformed glioma cells from copy number variants estimated by single-cell RNA-seq (scRNA-seq) (26). Here, we used SCOPE-seq to profile PpIX fluorescence and identify cell types at single-cell resolution in glioma resections from patients treated with 5-ALA, demonstrating that 5-ALA does not specifically label transformed glioma cells. We then used cell lines to examine potential 5-ALA labeling mechanisms through both metabolic conversion and cell-cell interactions. Experiments in animal models suggest that 5-ALA preferentially labels tumor tissue over normal brain and that this labeling contrast does not rely on BBB disruption.

## Results

### Integrated measurement of single cell PpIX fluorescence and expression profiles with SCOPE-seq2

To investigate whether 5-ALA specifically induces PpIX fluorescence in tumor cells, we used SCOPE-seq2 (29) to perform integrated fluorescence imaging and RNA-seq of individual cells collected from four surgically resected GBM specimens from patients (Fig. 1, Table S1) treated with 5-ALA (Fig. 2). We characterized morphological phenotypes and PpIX fluorescence of single cells captured in a microwell array device using a scanning epifluorescence microscope (Fig. S1). In SCOPE-seq2, we capture mRNA from individual cells using beads coated in barcoded DNA primers. We can identify the barcode sequence associated with each bead-cell pair in our device using sequential fluorescent probe hybridization. Because this same barcode sequence is associated with each single-cell cDNA library during Illumina sequencing, we can directly link PpIX fluorescence imaging data from each cell with its gene expression profile (see “Methods”).

**Figure 1.**
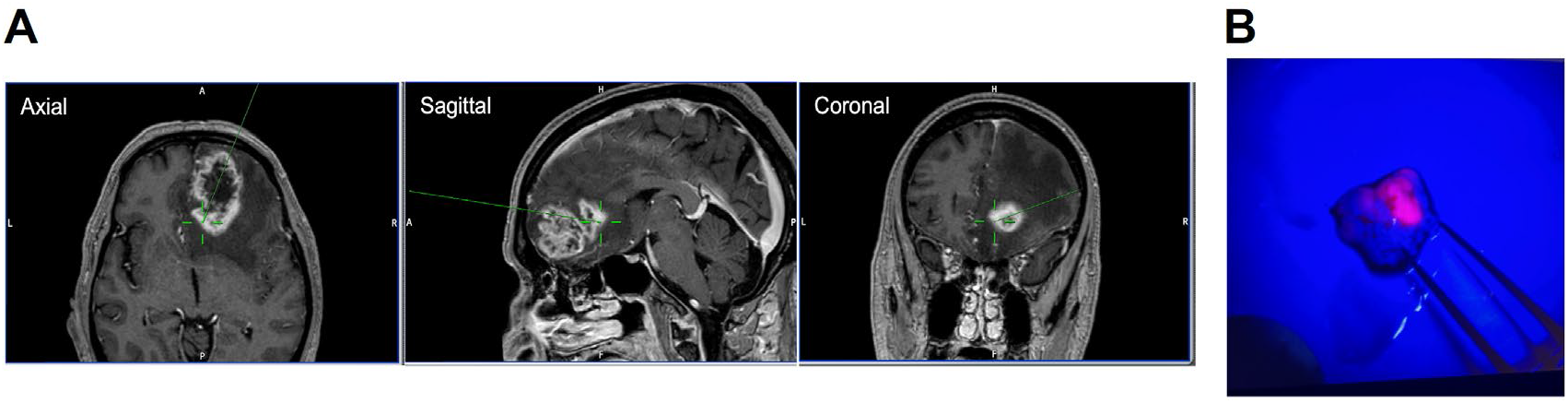
Representative MRI and intraoperative images of 5-ALA in GBM. (A) Pre-operative MRI annotated with intra-operative neuro-navigation. The position of resected tissue is indicated by the green crosshairs. (B)5-ALA induced PpIX fluorescence of resected tissue located at the tumor margin under violet-blue light with both 5-ALA positive (red fluorescence) and negative (blue) regions.

**Figure 2.**
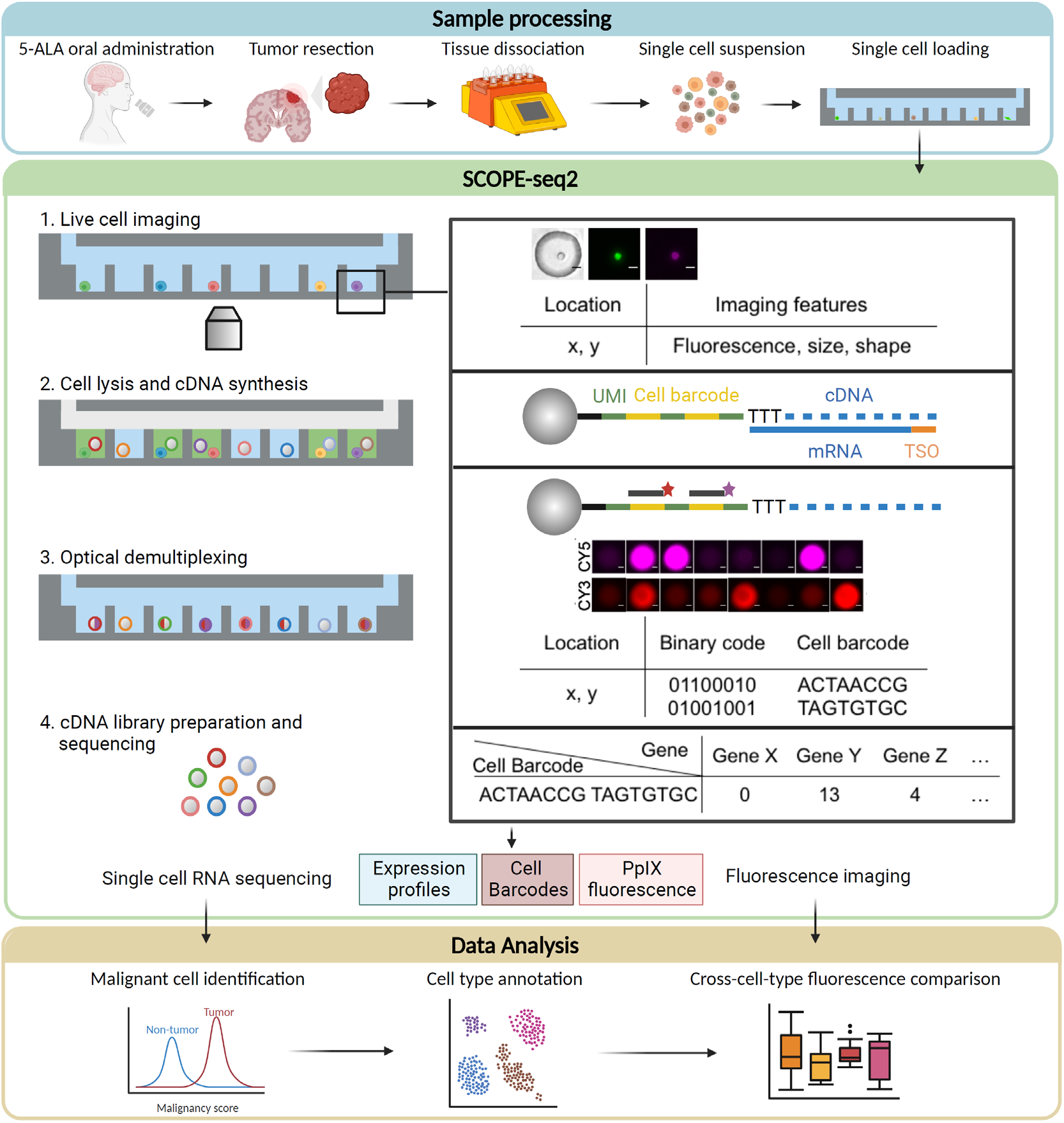
Schematic illustration of experimental and analytical methods using SCOPE-seq2 for comparing 5-ALA induced PpIX fluorescence among cell subpopulations in tumor microenvironment.

### 5-ALA labels both transformed glioma cells and untransformed cells in GBM tumor microenvironment

We obtained GBM surgical specimens from four patients who received 5-ALA pre-operatively (Table S1). For each patient, we collected one tissue specimen with intra-operatively observed PpIX fluorescence and performed SCOPE-seq2 including single cell fluorescence imaging and RNA-seq. For one of the cases (0826), we received a second specimen from a non-fluorescent region of the tumor and performed single cell fluorescence imaging in our device as a negative control. As expected, we observed that all four specimens for which PpIX fluorescence was observed intra-operatively exhibited higher PpIX fluorescence (Mann-Whitney U-test, p<0.01) by single-cell imaging than the specimen where PpIX fluorescence was not observed intra-operatively (Fig. 3A).

**Figure 3.**
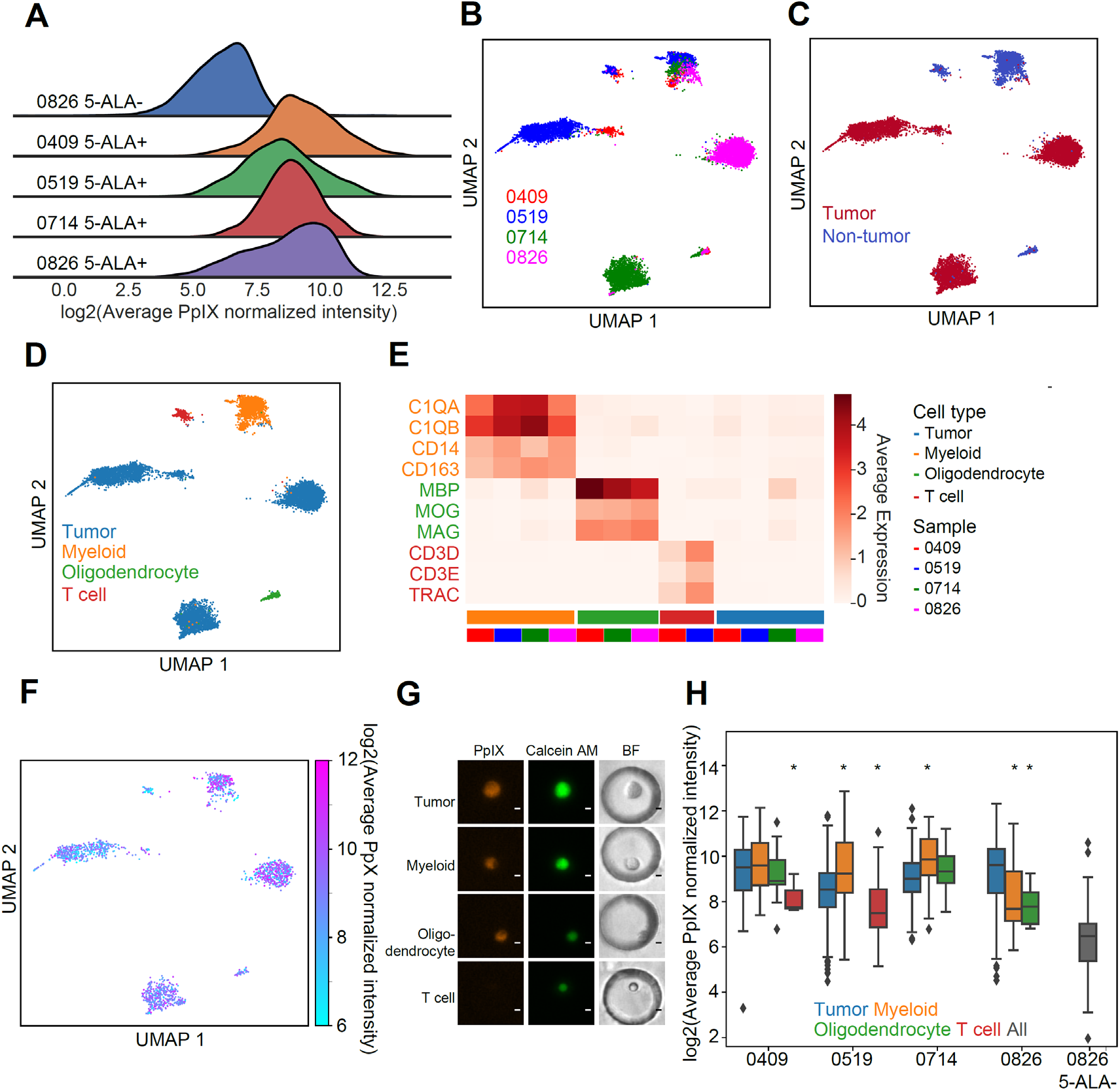
5-ALA labeling in tumor microenvironment. (A)Ridge plot of cell average PpIX fluorescence. (B)UMAP embedding of scRNA-seq expression profiles colored by patients. (C)UMAP embedding of scRNA-seq expression profiles colored by ploidy score. (D)UMAP embedding of scRNA-seq expression profiles colored by cell subpopulations. (E)Heatmap of average expression of lineage marker genes from cell types in the tumor microenvironment in each cell subpopulation and patient. (F)UMAP embedding of scRNA-seq expression profiles colored by average PpIX fluorescence. (G)Representative fluorescence images of each cell subpopulation in patient 0409. Scale bars: 5 μm. (H)Box plot of average PpIX fluorescence in each cell subpopulation and patient sample. PpIX fluorescence is compared between transformed glioma cells and other untransformed cell subpopulations of the same patient. Mann-Whitney U-test, *, q<0.05.

To investigate whether 5-ALA specifically labels tumor cells in tissue with visible PpIX fluorescence, we first identified cellular subpopulations from single-cell expression profiles. We performed unsupervised clustering and embedded the expression profiles in two dimensions using Uniform Manifold Approximation and Projection (UMAP, Fig. 3B-D) (30). We used aneuploidies, which can be inferred from scRNA-seq data, to identify malignantly transformed tumor cells as described previously (26). Chromosome 7 amplification and chromosome 10 deletion are two common aneuploidies in IDH wildtype GBM, and Chromosome 7 amplification may occur in IDH mutant high-grade glioma (26, 31). We verified the presence of these alterations using bulk, low-coverage whole genome sequencing (WGS) of the samples processed with SCOPE-seq2 (Fig. S2). We distinguished malignantly transformed tumor cells from non-malignant normal cells by identifying which cells harbored these copy number variants (CNVs, Fig. 3C, S3) from scRNA-seq. For example, cells with amplification of chromosome 7 express genes on chromosome 7 at higher levels on average. By examining the expression of cell lineage marker genes in each cluster, the non-malignant cells were further classified into myeloid cells (C1QA, C1QB, CD14, CD163), oligodendrocytes (MBP, MOG, MAG) and T cells (CD3D, CD3E, TRAC) (Fig. 3D, 3E). Overall, the cellular composition of the GBM specimens profiled in this study are highly concordant with previous analyses of GBM by scRNA-seq (26, 27).

To compare 5-ALA labeling among cell subpopulations, we analyzed the paired cell PpIX fluorescence and expression profiling data (Fig. 3F-H). We observed that all cell types in the tissue with visible PpIX fluorescence have higher PpIX fluorescence compared to cells from tissue without visible PpIX fluorescence (Mann-Whitney U-test, q<0.001, Fig. 3H, Table S2), suggesting that 5-ALA treatment results PpIX fluorescence in both transformed and untransformed cells within tumor microenvironment. In some cases, the transformed cells are not even the most fluorescent cells in the tumor. While T cells often have lower PpIX fluorescence than transformed tumor cells, oligodendrocytes are comparable and myeloid cells are brighter than transformed cells (Mann-Whitney U-test, q<0.05, Fig. 3H, Table S2).

To examine whether gene expression is related to cell PpIX fluorescence, we correlated the expression of protein coding genes with PpIX fluorescence in transformed cells for each patient. We ranked genes by their correlation and performed gene set enrichment analysis (GSEA) (32). This analysis did not identify any consistent, significant gene ontologies that were correlated with fluorescence intensity among the transformed glioma cells across patients (Fig. S4).

### Potential mechanisms of 5-ALA labeling of untransformed cells

We found that untransformed cells within GBM tumor microenvironment are fluorescently labeled after 5-ALA treatment, but the labeling mechanism of untransformed cells in tumor microenvironment is unknown. One potential labeling mechanism is that untransformed cells can uptake exogeneous 5-ALA and convert it into fluorescent PpIX. Since transformed cells can generate and secrete PpIX into the extracellular space after 5-ALA treatment (33, 34), another potential labeling mechanism is that untransformed cells uptake PpIX secreted by transformed cells. As shown above, myeloid cells such as brain-resident microglia are highly fluorescent in GBM after 5-ALA treatment (35). Thus, to determine whether untransformed cells can uptake 5-ALA and convert it into PpIX directly like transformed cells, we measured PpIX fluorescence of 5-ALA treated human microglial cell line HMC3 and human GBM neurosphere cell line TS543. We observed that both transformed GBM cells and untransformed myeloid cells have significantly increased PpIX fluorescence after 5-ALA treatment (t-test, p<0.001, Fig. 4A, 4B, Table S3). To determine whether cells can uptake PpIX secreted by transformed cells, we measured PpIX fluorescence of untreated TS543 or HMC3 cells after incubation with 5-ALA treated TS543 cells. To differentiate untreated cells from 5-ALA treated cells in images, we labeled untreated cells with live stain Calcein AM before incubation. To ensure that the observed PpIX fluorescence in 5-ALA untreated cells is not induced by 5-ALA directly, we washed 5-ALA treated cells thoroughly with PBS to remove exogeneous 5-ALA and resuspended the treated cells in fresh medium before incubation with 5-ALA untreated cells, and we also measured the PpIX fluorescence of 5-ALA untreated cells incubated with the resuspension supernatant. We found that both GBM cells and untransformed microglia have significantly increased PpIX fluorescence after incubation with 5-ALA labeled GBM cells (t-test, p<0.001, Table S4) but not after incubation with the supernatant (Fig. 4C, 4D). These results suggest that PpIX fluorescence of untransformed cells in GBM tumor microenvironment can be induced directly by 5-ALA uptake and conversion to PpIX, or indirectly by uptake of PpIX secreted from surrounding transformed cells.

**Figure 4.**
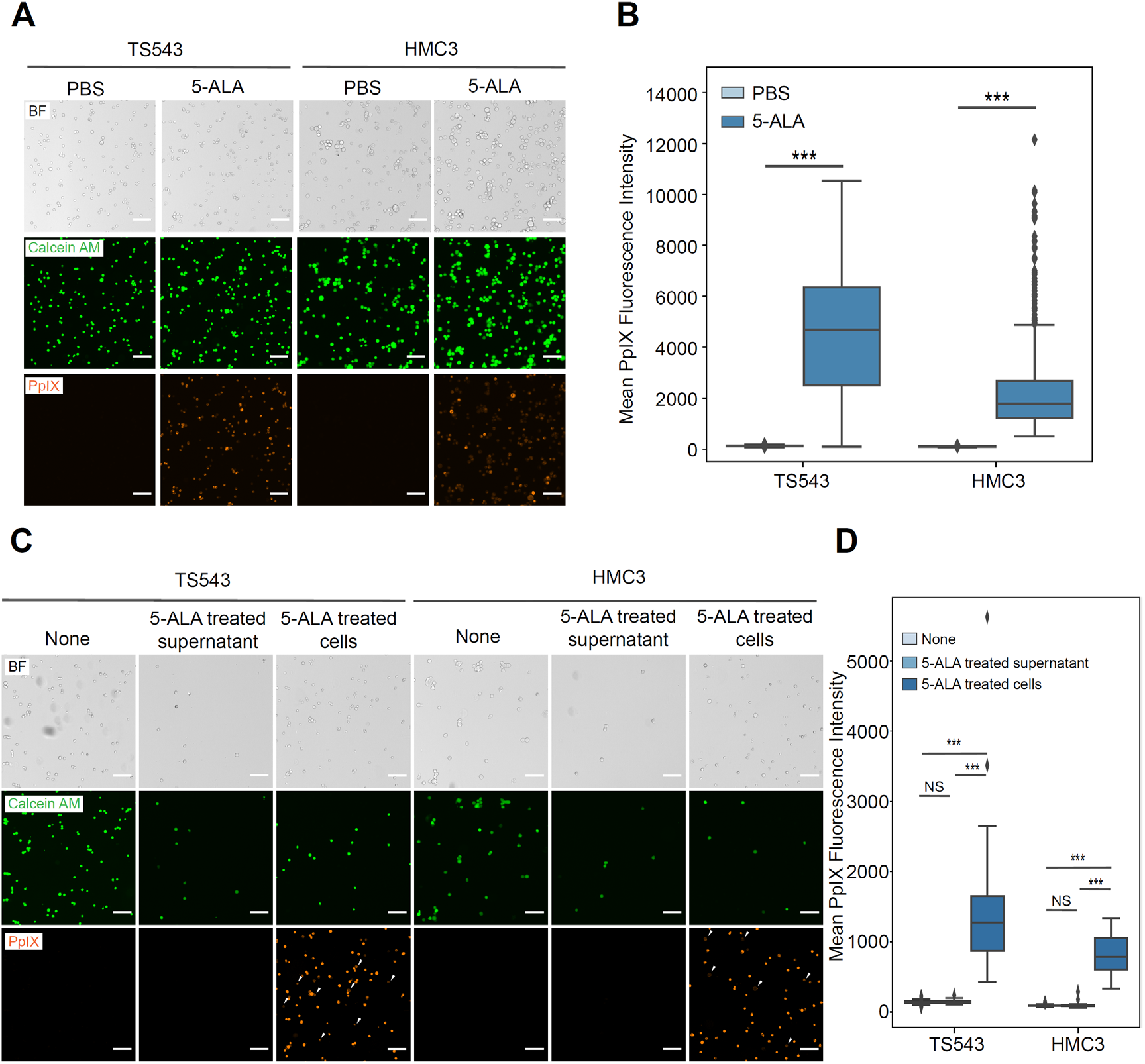
Potential mechanisms of 5-ALA labeling of untransformed cells. (A)Example images of TS543 (glioma) and HMC3 (cultured microglia) cells treated with PBS control or 5-ALA. BF, bright field; Calcein AM, live cells; PpIX, 5-ALA labeling. Scale bars: 100 μm. (B)Box plot shows the quantification of PpIX fluorescence of TS543 and HMC3 cells treated with PBS control or 5-ALA. Unpaired t-test, ***, p<0.001. (C)Representative images of 5-ALA untreated TS543 and HMC3 cells before (None) or after treatment with 5-ALA treated cell supernatant or 5-ALA treated TS543 cells. BF, bright field; Calcein AM, 5-ALA untreated cells; PpIX, 5-ALA labeling. White arrows, 5-ALA untreated cells. Scale bars: 100 μm. (D)Box plot shows the quantification of PpIX fluorescence of 5-ALA untreated TS543 and HMC3 cells before (None) or after treatment with 5-ALA treated cell supernatant or 5-ALA treated TS543 cells. Unpaired t-test, ***, p<0.001; NS, not significant.

### 5-ALA preferentially labels tumor tissue over non-neoplastic brain tissue regardless of blood-brain-barrier status

Although 5-ALA labels both transformed and untransformed cells in GBM, 5-ALA is reported to preferentially label tumor tissue (9). While this preference could arise from specific labeling of transformed glioma cells, our SCOPE-seq2 experiments eliminate this possibility. Another possibility is that tumor tissue-selective labeling is due to blood-brain-barrier (BBB) disruption by the tumor, making 5-ALA more accessible to tumor tissue compared with non-neoplastic brain tissue. A third possibility is that metabolic alterations in the GBM microenvironment could results in PpIX accumulation. To remove the effect of BBB during 5-ALA labeling, we generated acute brain slice cultures from mice injected with GL261 cells, resulting in the formation of syngeneic brain tumors, and from wild type mice as described previously (36) and treated the slice cultures with 5-ALA *ex vivo* (Fig. 5A). After 5-ALA treatment, we generated thin sections for each slice and measured the tissue average PpIX fluorescence using confocal microscopy. We then used H&E-stained adjacent sections to differentiate tumor and normal tissues. We observed that tumor tissues in GL261 mice show significantly increased average PpIX fluorescence across the imaging field after 5-ALA treatment (t-test, p<0.001, Fig 5B, 5C, Table S5), while the average PpIX fluorescence of normal brain tissues in both GL261 and wild type mice do not differ significantly. We also observed that tumor tissues have higher average PpIX fluorescence than normal tissues after 5-ALA treatment in GL261 mice (t-test, p<0.001), indicating that 5-ALA selectively labels GBM tumor tissue at the bulk level even given uniform access to the tumor and normal brain. These results suggest that although 5-ALA labeling is not specific to transformed glioma cells, selective labeling of tumor tissue over non-neoplastic brain tissue is independent of BBB disruption.

**Figure 5.**
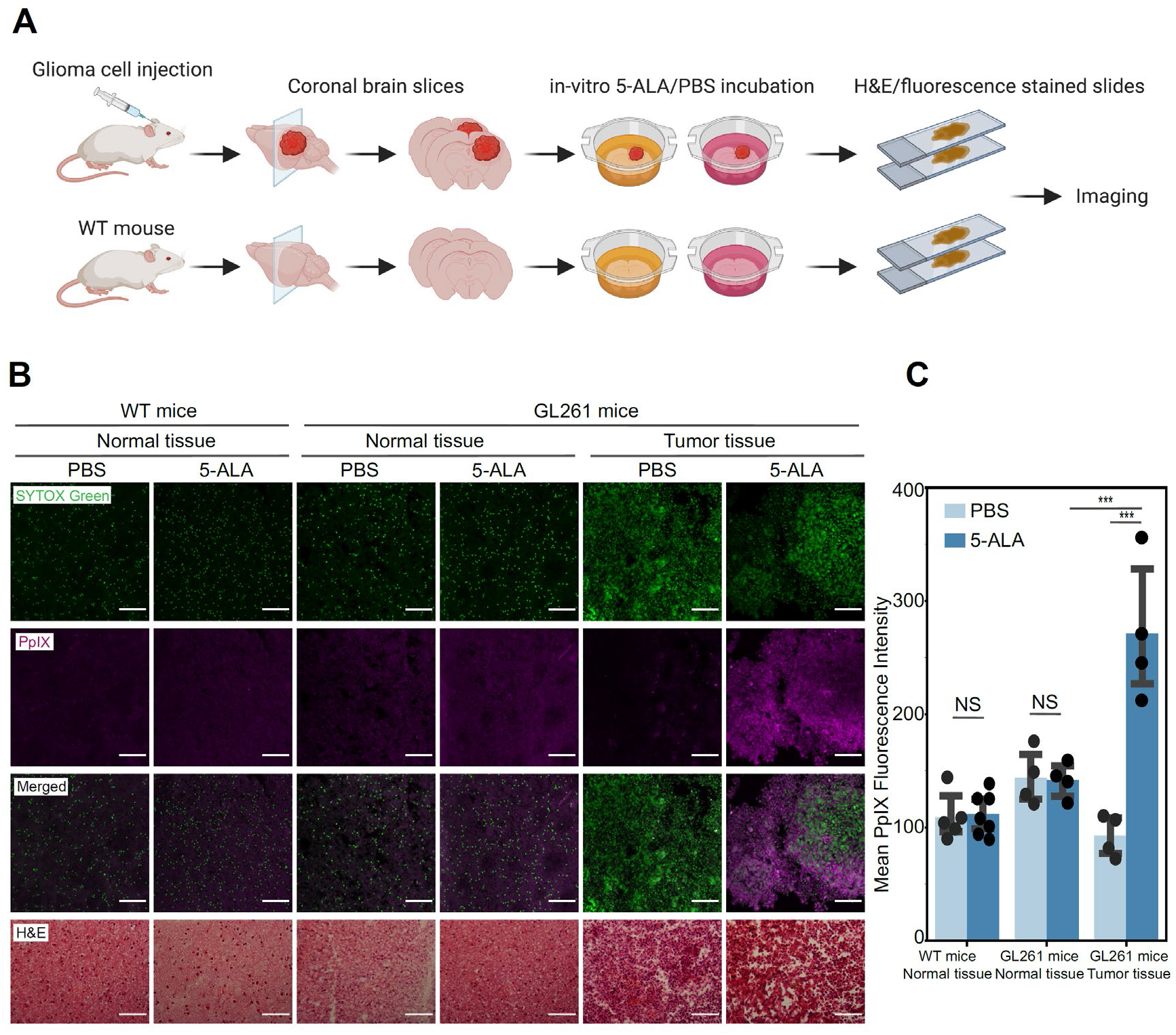
In vitro treatment of 5-ALA on mouse slice cultures. (A)Schematic illustration of in vitro 5-ALA treatment of mouse brain slice cultures and imaging. (B)Representative images of normal and tumor tissues in wild type and GL261 mouse brain slice cultures treated with PBS control or 5-ALA. SYTOX Green, nuclei; PpIX, 5-ALA labeling; H&E, hematoxylin and eosin. Scale bars: 100 μm. (C)Box plot shows the quantification of the average PpIX fluorescence of normal and tumor tissues treated with PBS control or 5-ALA. Each dot represents a slice culture. Unpaired t-test, ***, p<0.001; NS, not significant.

## Discussion

In this study, we sought to determine the specificity of fluorogenic labeling by 5-ALA in high-grade glioma patients with single-cell resolution. While prior studies have reported the predictive value of 5-ALA labeling for identifying tumor tissue (7, 9, 18-20), the mechanism underlying this apparent specificity is unclear. One possibility is that transformed glioma cells themselves are specifically labeled by 5-ALA due to unique metabolic properties. Another is that the tumor microenvironment as a whole shares metabolic alterations that are conducive to labeling or that certain cells specifically metabolize 5-ALA and others take up the resulting PpIX secreted by these cells. Less sophisticated FGS agents like fluorescein likely just label the extracellular space in regions of BBB disruption. Thus, although 5-ALA is clearly producing intracellular PpIX, BBB breakdown could also facilitate specific delivery of 5-ALA to glioma tissue. However, our results with the GL261 slice cultures show that selective labeling of tumor tissue is not dependent on BBB breakdown.

By combining scRNA-seq and live cell imaging with SCOPE-seq2, we learned that many cell types, including untransformed cells in the glioma microenvironment, are brightly labeled by PpIX in glioma surgical specimens from patients who received 5-ALA. In fact, 5-ALA treatment does not even preferentially label transformed glioma cells in some tumors, and myeloid cells can be significantly brighter. Nonetheless, our studies in acute slice cultures of a glioma mouse model and normal brain tissue showed that glioma tissue is specifically labeled by 5-ALA. In this experimental paradigm, differences in BBB status between tumor tissue and normal brain are irrelevant, because we administer 5-ALA directly to tissue slices. Further experiments in cultured human microglia and patient-derived glioma neurospheres showed that both cell types are capable of converting 5-ALA into PpIX and that PpIX can be secreted and taken up by unlabeled cells. These results suggest that the apparent specificity of 5-ALA for glioma tissue is not due to BBB disruption and that both transformed glioma cells and cells in the microenvironment can likely metabolize 5-ALA into PpIX. Furthermore, glioma cells can likely secrete PpIX, which can then be taken up by both unlabeled glioma cells and microglia.

Previous studies have reported that the metabolic status of a cell could affect intracellular PpIX accumulation (8, 11-14), thus we might have expected to see consistent enrichment of metabolic pathways associated with PpIX fluorescence intensity in transformed glioma cells. One possible reason that we do not find any consistently enriched metabolic pathway in our analysis is that there are multiple possible mechanisms by which a tumor cell could become labeled, including uptake of PpIX secreted by other cells in the microenvironment.

While previous studies have reported the utility of 5-ALA in FGS (7, 9), our results demonstrate that the PpIX fluorescence arising from 5-ALA treatment does not originate exclusively from transformed glioma cells. Furthermore, although 5-ALA is typically considered to be an intracellular labeling method, there could be substantial extracellular PpIX secreted by multiple cell types in and around a tumor. This raises the possibility that 5-ALA could spuriously label untransformed cells in the glioma margins, where labeling specificity is most critical. While 5-ALA likely still offers specificity advantages over simpler FGS agents like fluorescein, future studies should systematically evaluate the performance of these two labeling approaches, preferably by simultaneous treatment of the same patients.

## Methods

### Human clinical sample preparation

Patients were given 5-ALA (Gleolan) at 20mg/kg approximately 4 h before surgery. Tissue specimens were stratified into 5-ALA positive (red) and negative (blue) tissues by the surgeon under violet-blue light and collected immediately after surgical removal. Collected tissue samples were dissociated using the Adult Brain Dissociation kit (Miltenyi Biotec, cat # 130-107-677) on gentleMACS Octo Dissociator with Heaters (Miltenyi Biotec) according to the manufacturer’s instructions with following modifications. Calcein AM (ThermoFisher Scientific, cat# C3100MP) was added to the dissociation enzyme mix to a final concentration of 4µM to label live cells. Cells were finally resuspended in 1x TBS buffer at a concentration of 1000 cells per μl.

### SCOPE-seq2 imaging and scRNA-seq library preparation and sequencing

Dissociated cells were profiled using SCOPE-seq2 method as previously described (29). A microwell array device was filled with wash buffer (20 mM Tris–HCl pH7.9, 50 mM NaCl, 0.1% Tween-20), stored in a humid chamber one day before use and flushed with 1x TBS before cell loading. A single cell suspension was loaded into the microwell device, and untrapped cells were flushed out with 1x TBS. The device was scanned under an inverted automated microscope (Nikon Eclipse Ti2) in two fluorescence channels and the bright field channel with wide-field 10 × 0.3 NA objective (Nikon, cat# MRH00101) in ∼ 10 minutes. Bright-field images were taken using an RGB light source (Lumencor, Lida). Fluorescence images were taken using LED light source (Lumencor, spectra x). Calcein AM fluorescence was excited at 470nm and emission collected at 500-550nm. PpIX fluorescence was excited at 395 nm and emission collected at 600-660nm. Optically barcoded (OBC) beads (Chemgenes) were then loaded to the microwell device, and untrapped beads were flushed out with 1x TBS.

The microwell device containing cells and beads was connected to the computer-controlled reagent and temperature delivery system for single cell RNA sequencing (scRNA-seq). The microwells were in lysis buffer (1% 2-mercaptoethanol (Fisher Scientific, cat# BP176-100), 99% Buffer TCL (Qiagen, cat# 1031576), sealed with perfluorinated oil (Sigma-Aldrich, cat# F3556-25ML) at 50 °C for 20 min to promote cell lysis, and then at 25 °C for 90 min for mRNA capture. Wash buffer supplemented with RNase inhibitor (0.02 U/µL SUPERaseIN (Thermo Fisher Scientific, cat# AM2696) in wash buffer) was flushed through the device to unseal the microwells. Reverse transcription mixture (1X Maxima RT buffer, 1 mM dNTPs, 1 U/µL SUPERaseIN, 2.5 µM template switch oligo, 10 U/µL Maxima H Minus reverse transcriptase (Thermo Fisher Scientific, cat# EP0752), 0.1% Tween-20) was flowed into the device followed by an incubation at 25 °C for 30 min and then at 42 °C for 90 min. Exonuclease I reaction mixture (1X Exo-I buffer, 1 U/µL Exo-I (New England Biolabs, cat# M0293L)) was pipetted into the device followed by an incubation at 37 °C for 45 min. TE/TW buffer (10 mM Tris pH 8.0, 1 mM EDTA, 0.01% Tween-20) was flushed through the device. Exonuclease I reaction mixture (1X Exo-I buffer, 1 U/µL Exo-I (New England Biolabs, cat# M0293L)) was pipetted into the device followed by an incubation at 37 °C for 45 min. The device was then washed with TE/TW buffer (10 mM Tris pH 8.0, 1 mM EDTA, 0.01% Tween-20).

The microwell device containing the beads with cDNAs was connected to a computer-controlled reagent delivery and scanning system for optical demultiplexing. Eight-cycle probe hybridization and stripping were performed to identify cell barcode sequence corresponded to imaging positions. Probes were stripped with melting buffer (150 mM NaOH) for 10 min, then washed with imaging buffer (2xSSC, 0.1% Tween-20). Background device scan images were taken in the bright-field, Cy3 (555nm excitation) and Cy5 (649 excitation) channels with Quad band filter set (Chroma, cat# 89402) to ensure probe strip efficiency. Probes were hybridized with hybridization solution (imaging buffer supplemented with a pool of Cy3 or Cy5 fluorescent labeled probes that were reverse complemented to the cell barcode sequences) for 10 min, then washed with imaging buffer. Probe hybridization images were taken in the bright-field, Cy3 and Cy5 channels.

After optical demultiplexing, microwells were sealed with perfluorinated oil, and then the device was cut into 10 regions. Beads from each region were extracted separately, washed sequentially with TE/SDS buffer (10 mM Tris–HCl, 1 mM EDTA, 0.5% SDS), TE/TW buffer, and nuclease-free water. cDNA amplification was performed in 50 µL PCR solution [1X Hifi Hot Start Ready mix (Kapa Biosystems, cat# KK2601), 1 µM SMRTpcr primer (Supplementary Table S6)], with 16 amplification cycles (95 °C 3 min, 4 cycles of (98 °C 20 s, 65 °C 45 s, 72 °C 3 min), 12 cycles of (98 °C 20 s, 67 °C 20 s, 72 °C 3 min), 72 °C 5 min). PCR product was purified using SPRI paramagnetic bead (Beckman, cat# A63881) with a bead-to-sample volume ratio of 0.6:1. cDNA library preparation was performed using illumina Nextera library perp kit (Illumina, FC-131-1024) with 0.6ng cDNA as input. The scRNA-seq libraries were pooled and sequenced on an Illumina NextSeq 500 sequencer using a 75 cycle High Output Kit (26-cycle read 1, 58-cycle read 2, and 8-cycle index 1) with custom read 1 primer.

### SCOPE-seq2 imaging and sequencing data analysis

Live cell fluorescence images were analyzed using ImageJ as previously described (29). Microwells with cells were identified and analyzed individually within the smallest bounding square of the corresponding microwell. Cell regions-of-interest (ROIs) were identified from the Calcein AM fluorescence, and the remaining area was identified as background. Then the cell average PpIX fluorescence were measured and normalized by subtracting the background PpIX fluorescence. Cell multiplets identified as multiple ROIs in one microwell or manually were removed from downstream analysis.

To identify cell barcode sequences, bead ROIs were identified in the bright field channel, then bead fluorescence intensity measurements were performed for two channels (Cy3 and Cy5). Measurements of the same bead is registered by the nearest bead positions across 8-cycle images. Cell barcode calling was performed using a bead-by-bead algorithm as previously described (29). To associate cell barcodes with cell imaging phenotypes, microwell positions in optical demultiplexing images were mapped to the positions in live cell fluorescence images.

To analyze the scRNA-seq data from SCOPE-seq2, we first extracted the cell-identifying barcode and UMI from Read 1 based on the designed oligonucleotide sequence, NN(8-nt Cell Barcode S)NN(8-nt Cell Barcode Q)NNNN. The 192 8-nt cell barcode sequences have a Hamming distance of at least three for all sequence pairs. Therefore, we corrected at most one substitution error in the cell barcode sequences. We only keep reads with a complete cell barcode. Next, we align the reads from Read 2 to a merged human genome (GRCh38) using STAR v.2.7.0 aligner (37) after removing 3′ poly(A) tails (indicated by tracts of > 7 A’s) and fragments with fewer than 24 nucleotides after poly(A) tail removal. Only reads that were uniquely mapped to exons on the annotated strand were included for the downstream analysis. Reads with the same cell barcode, UMI (after one substitution error correction) and gene mapping were considered to originate from the same cDNA molecule and collapsed. Finally, we used this information to generate a molecular count matrix.

Imaging and the expression data of the same single cell were then linked by the same cell barcode sequence.

### scRNA-seq clustering and visualization

For clustering and visualization of the merged dataset from four samples, we first subsampled the merged dataset so that every sample has the same number of cells. We then factorized the subsampled gene count matrix using the single-cell hierarchical Poisson factorization (scHPF) algorithm (38) with default parameters and k = 17, and identified top 100 genes in each factor. We removed nuisance factors associated with heat shock protein genes and cell stress. Clustering and visualization were performed on the complete merged dataset. To cluster the expression profiles, we computed Pearson correlation distance from variable-gene count matrix, constructed a k-nearest neighbors’ graph with k=20 and clustered using the Phenograph implementation of Louvain community detection (39). To visualize the expression profiles, we generated a two-dimensional embedding of the Pearson correlation distance matrix using Uniform Manifold Approximation and Projection (UMAP) (30).

For clustering and visualization of expression profiles from each individual sample, variable genes were selected as genes detected in fewer cells than expected given their apparent expression level using the drop-out curve method as previously described (26). Phenograph clustering and UMAP embedding were processed as described above.

### Whole genome sequencing

Genomic DNA was extracted from each bulk tumor tissue and PBMC sample using the DNeasy Blood & Tissue Kits (Qiagen) according to the manufacturer’s instructions. DNA library was prepared using Nextera DNA library prep kit according to the manufacturer’s instructions. Raw sequencing data were aligned to the human genome (GRCh38) using bwa mem (40) and analyzed as described previously (26). Briefly, we computed the number of de-duplicated reads that aligned to each chromosome for each patient and divided this by the average number of de-duplicated reads that aligned to each chromosome for two diploid germline samples after normalizing both by total reads. We then normalized this ratio by the median ratio across all somatic chromosomes and multiplied by two to estimate the average copy number of each chromosome.

### Transformed tumor cell identification

The cell aneuploidy analysis was performed based on the scHPF model as described previously (36). For each sample, gene count matrix was factorized using scHPF method with default parameters and k = 15. To compute the scHPF-imputed expression matrix, we multiplied the gene and cell weight matrix (expectation matrix of variable *θ* and *β* in the scHPF model and then log-transformed the result matrix as log_2_ (expected counts / 10000 + 1). For IDH wildtype GBM tumors, we defined a ploidy score as the difference between the average expression of Chr. 7 genes to that of Chr. 10 genes, < log_2_(Chr. 7 Expression) > − < log_2_(Chr. 10 Expression) >; for IDH mutant astrocytoma samples, the ploidy score is defined as the average expression of Chr. 7 genes, < log_2_(Chr. 7 Expression) >. We fitted a double Gaussian distribution to the ploidy score and a threshold at 1.96 standard deviations (95% confidence interval) below the mean of the Gaussian with the higher mean is defined as threshold to identify tumor and non-tumor cells. Cells with ploidy scores lower than the threshold in the tumor cell Phenograph clusters and cells with ploidy scores higher than the threshold in the non-tumor cell Phenograph clusters were removed from the downstream analysis.

### Gene set enrichment analysis

We did the following analysis on transformed cells for each individual patient. We computed Spearman’s correlation using *spearmanr* command in the Python module *scipy* between the normalized PpIX fluorescence intensity and the log2(counts per million+1) normalized expression levels for each protein-coding gene detected in at least 10% of cells. The resulting p-values were corrected using the Benjamini–Hochberg method as implemented by the *multipletests* function in the Python package *statsmodels*. We then ranked all of the genes by (Spearman correlation coefficient) * (-log10(q value)) and used this as input for the gene set enrichment analysis (GSEA). We used gene sets from the Molecular Signatures Database C5 collection of gene ontologies and genes detected in less than 10% cells were removed from the gene sets. We performed GSEA with the pre-ranked gene list and the filtered gene sets in the pre-ranked mode with “classic” enrichment statistic.

### Cell lines

TS543 human glioma cell line was cultured in human NeuroCult™ NS-A Basal Medium (Stem Cell Technologies, cat #05751), supplemented with 10% NeuroCult™ Proliferation Supplement, heparin (2 μg/ml, Stem Cell Technologies, cat# 7980), human bFGF (10 ng/ml, Stem Cell Technologies, cat# 78003) and human EGF (20 ng/ml, Stem Cell Technologies, cat# 78006). HMC3 human microglia cell line is cultured in Eagle’s Minimum Essential Medium (EMEM, ATCC, cat# 30-2003) supplemented with 10% fetal bovine serum (FBS, Gibco, cat# A3160402). Cell lines are incubated at 37 °C and 5% CO_2_.

For 5-ALA treatment, 5-ALA HCL (sigma, cat# A7793) was dissolved in 1x PBS and pH is adjusted to 7.4 using NaOH. Cells were incubated in medium with 1 mM 5-ALA for 4 hours at 37 °C and 5% CO2. TS543 or HMC3 cells were then dissociated with TrypLE Express Enzyme (Gibco, cat# 12605028), stained with Calcein AM live stain dye (ThermoFisher Scientific, cat# C3100MP) at 4 µM for 5 minutes.

For incubation with 5-ALA treated cells, we first treated human glioma TS543 cells with 5-ALA as described above. Dissociated 5-ALA treated TS543 cells were washed with 1x PBS for three times and resuspended in fresh TS543 culture medium without exogenous 5-ALA. We split the resuspension into two parts, one was referred to as 5-ALA treated cells, the other was further centrifuged and the supernatant was referred to as 5-ALA treated supernatant. 5-ALA untreated TS543 or HMC3 cells were dissociated with TrypLE Express Enzyme and stained with Calcein AM as described above. We then mixed 5-ALA untreated cells with treated cells and incubated them at 37 °C and 5% CO2 for 1 hour.

### Mouse tissue preparation

GL261 cells were grown in DMEM-10% FBS, at 37°C with 5% CO2. Adult C57Bl/6 mice were anesthetized with Ketamine-Xylazine (100 mg/kg and 10mg/kg, respectively), and assessed for lack of reflexes by toe pinch. Hair was shaved and scalp skin incised. After identifying bregma, a burr hole was made with a 17-gauge needle 2mm lateral and 2mm anterior to the bregma. Intracranial injection (5×103 cells in 1μL) was performed under stereotactic guidance, 2mm deep, using a Hamilton syringe at a flow rate of 0.3μL/min. Tumor growth was assessed through MRI. Mice were sacrificed at 25dpi by cervical dislocation. The brain was removed, placed into ice-cold PBS and cut into 300-500µm sections using a McIlwain Tissue Chopper. Sections were transferred onto Millicell cell culture inserts (0.4µm, 30mm diameter) that were placed in 6-well plates containing 1.5mL of medium consisting of DMEM/F12 with N-2 Supplement and 1% antimycotic/antibiotic. Slices were first rested overnight with the medium in a humidified incubator at 37°C and 5% CO2. Then, the medium was replaced with pre-warmed medium containing 1 mM 5-ALA or PBS as control. Slices were then cultured with the treatment medium in a humidified incubator at 37 °C and 5% CO2 for 4 h. After 5-ALA treatment, slices were fixed by 4% PFA (Thermo Scientific, cat# 28908) for 24 hours at 4 °C and incubated in 30% sucrose at 4 °C until they are embedded in OCT for cryosectioning into 16μm sections. Adjacent tissue slides of each slice culture were used for PpIX fluorescence measurement and H&E. For PpIX fluorescence measurement, slides were warmed up to room temperature, rehydrated with 1x PBS for 10 minutes, stained with 10 nM SYTOX Green for 5 minutes, washed with 1x PBS and mounted with Fluoromount-G mounting medium (Invitrogen, cat# 00-4958-02). For H&E staining, slides were placed in 1x PBS for 5 minutes, stained with Eosin and Hematoxylin (5-10 seconds each), washed with dH2O until clear, dehydrated (once in 70%, twice in 90%, twice in 100% Ethanol, twice in Xylene, 10 seconds each step) and mounted with Permount.

### Microscopy and imaging analysis

For cell cultures, images were taken with an inverted automated microscope (Nikon Eclipse Ti2) in two fluorescence channels and the bright field channel with wide-field 10 × 0.3 NA objective (Nikon, cat# MRH00101). Bright-field images were taken using an RGB light source (Lumencor, Lida). Fluorescence images were taken using LED light source (Lumencor, spectra x). Calcein AM fluorescence was excited at 470nm and emission collected at 500-550nm. PpIX fluorescence was excited at 395 nm and emission collected at 600-660nm. We used ImageJ for image analysis. Cell regions-of-interest (ROIs) were identified from the Calcein AM fluorescence using *threshold* and *particle analyzer*. Then the cell mean PpIX fluorescence was measured for each individual cell. To compare PpIX fluorescence across treatment conditions in cell lines, we first down-sampled cells each treatment condition into the same number of cells and then performed unpaired t-test as implemented as *ttest_ind* from Python module *scipy*.

For mice slice culture slides, fluorescence images were taken using a Nikon A1R MP confocal microscope with a 25 × 1.1 NA objective. SYTOX Green fluorescence was excited at 488 nm and emission collected at 500-550nm. PpIX fluorescence was excited at 403 nm and emission collected at 663-738nm. H&E images were taken using an upright microscope (Nikon Optiphot) with 20 × 0.3 NA objective. We used ImageJ for image analysis. Tissue regions were identified from the SYTOX Green fluorescence using *threshold*; regions with SYTOX Green intensity <10 was considered as region without tissue presence. Then the mean PpIX fluorescence of the tissue was measured for each slice culture. To compare PpIX fluorescence across treatment conditions in slice cultures, we performed unpaired t-test as implemented as *ttest_ind* from Python module *scipy*.

### Statistics

To compare PpIX fluorescence across cell types, we first down-sampled cells each cell type and each patient into the same number of cells. Then we performed the two-sided Mann-Whitney U-test as implemented by the *mannwhitneyu* command in the Python module *scipy* between two cell subpopulations within each patient sample. The resulting p values were corrected using the Benjamini–Hochberg method as implemented by the *multipletests* function in the Python package *statsmodels*.

### Study Approval

All of the animal protocols used in these studies were approved by the Columbia University Institutional Animal Care and Use Committee (IACUC). For studies of human specimens, all tissue was procured from de-identified patients who provided written informed consent through a protocol approved by the Columbia University Institutional Review Board (IRB-AAAJ6163).

## Acknowledgements

P.A.S. was funded by 75N910019C00029 from NIH/NCI. P.A.S., P.C., and J.N.B. were funded by R01NS103473 from NIH/NINDS. This research was funded in part through the NIH/NCI Cancer Center Support Grant P30CA013696 and used the Genomics and High Throughput Screening Shared Resource, Molecular Pathology Shared Resource, and the Confocal and Specialized Microscopy Shared Resource of the Herbert Irving Comprehensive Cancer Center at Columbia University.

## Author contributions

Z.L. and P.A.S. conceived the study and designed the experiments. Z.L. conducted the SCOPE-seq experiments. J.N.B. and P.C. procured the GBM tissue. Z.L., A.M., J.F., C.K., C.P.S. and N.H. performed mice tissue slice experiments. Z.L. analyzed the data with input from P.A.S. Z.L. and P.A.S. wrote the manuscript. All authors edited, read, and approved the final manuscript.

### Data availability

The sequencing data and count matrices reported in this paper are available in the following database: Gene Expression Omnibus GSE218331 (https://www.ncbi.nlm.nih.gov/geo/query/acc.cgi?acc=GSE218331). Code is available at: https://github.com/simslab/SCOPEseq2 and https://github.com/simslab/cluster_diffex2018.

