## Supplementary Figures 1-4 and Supplementary Table Legends for "Single-Cell Analysis of 5-ALA Intraoperative Labeling Specificity for Glioblastoma"

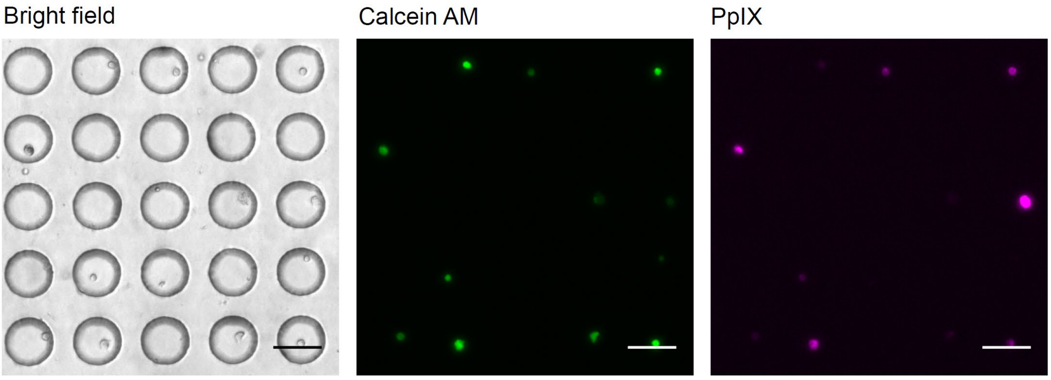


**Figure S1**. Representative fluorescence images of single cells from 5-ALA positively labeled tumor tissue captured in microwells. Scale bars: 50 μm.

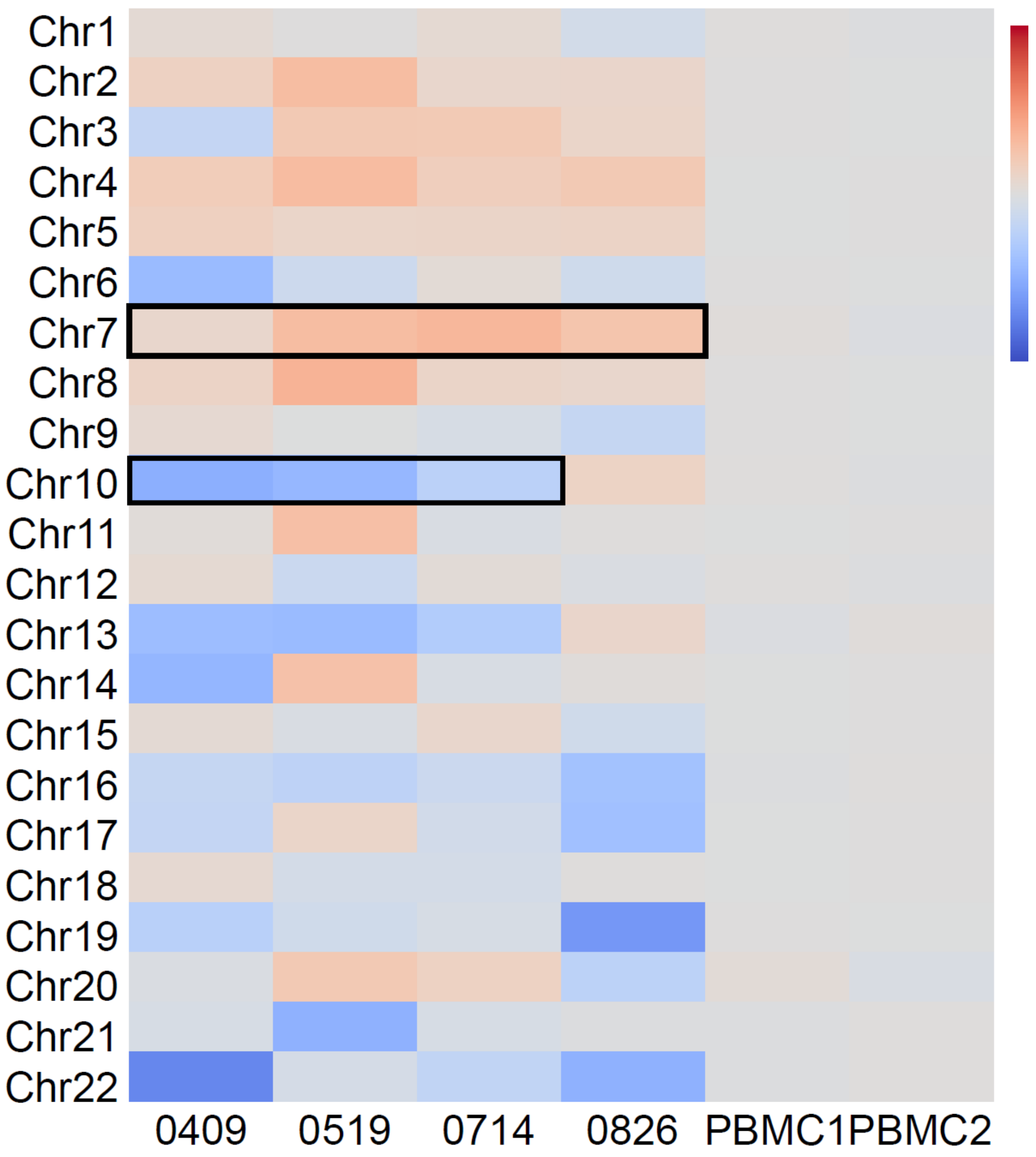


**Figure S2.** Heatmap of estimated chromosomal copy numbers from whole genome sequencing in GBM tumor tissue samples of 4 patients and 2 human peripheral blood mononuclear cell control samples.

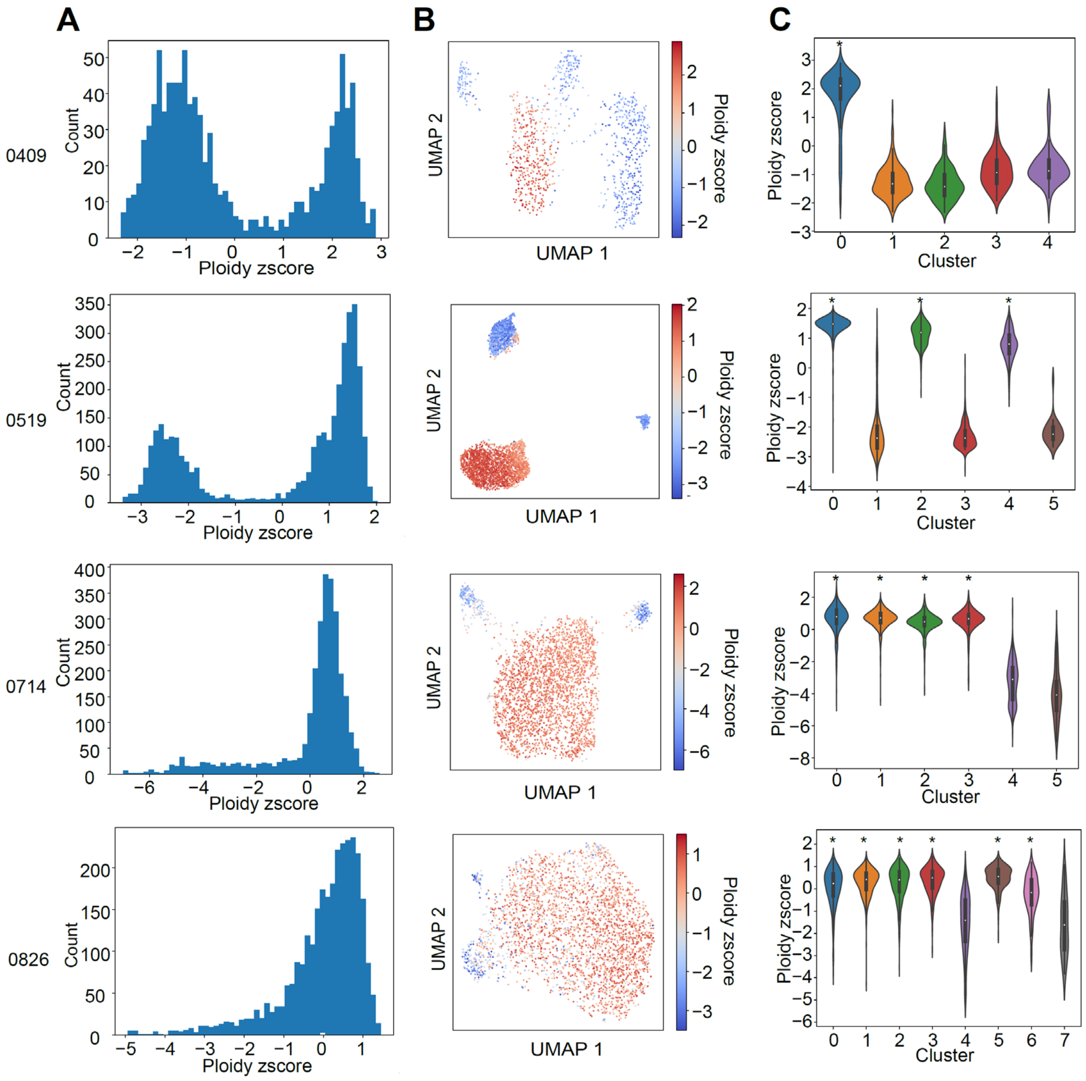


**Figure S3.** Identification of malignantly transformed tumor cells.

(A) Histograms of ploidy z-scores in each patient. Ploidy score is defined as log ratio of average Chr.7 to Chr.10 expression in IDH wildtype GBMs, and log-transformed average Chr.7 expression in the IDH mutant astrocytoma.

(B) UMAP embeddings of scRNA-seq expression profiles in each patient colored by ploidy z-scores.

(C) Violin plots of ploidy z-scores in each Phenograph cluster of expression profiles and patients. *, malignant transformed cell clusters with higher ploidy z-scores.

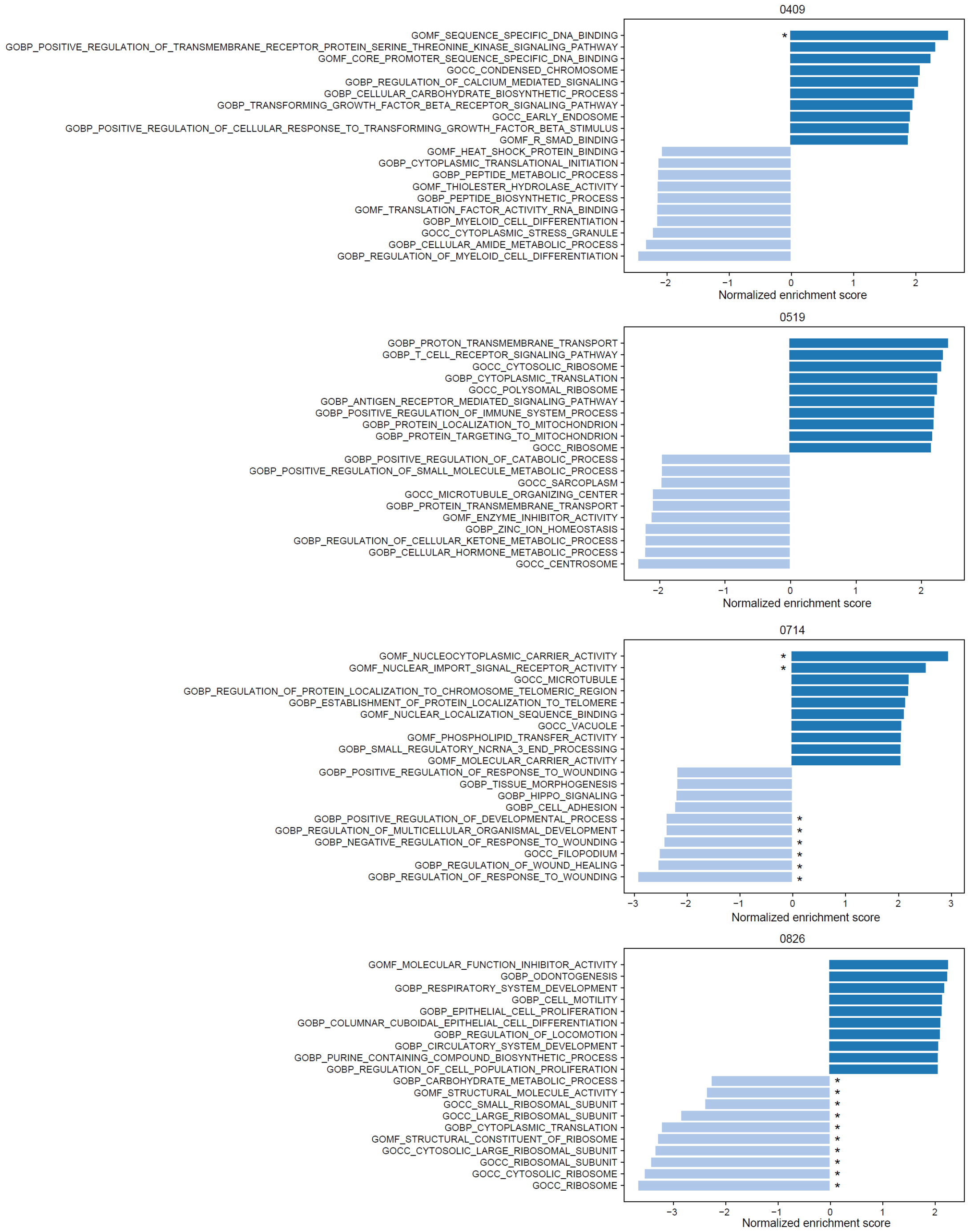


**Figure S4.** Normalized enrichment scores for top 10 gene ontology terms that are positively or negatively associated with the PpIX fluorescence. *, FDR<0.25.

### Supplementary Table Legends

**Table S1.** Summary of patients and tissue specimens.

**Table S2.** Mann-Whitney U-test of PpIX fluorescence among cell subpopulations.

**Table S3.** Unpaired t-test of PpIX fluorescence in cell cultures before and after 5-ALA treatment.

**Table S4.** Unpaired t-test of PpIX fluorescence in cell cultures before and after treatment with 5-ALA treated cells or supernatant.

**Table S5.** Unpaired t-test of PpIX fluorescence in mouse slice cultures before and after 5-ALA treatment.
